# Effects of nonpharmacological manipulations and repeated xanomeline treatment on methamphetamine-vs-food choice in Sprague Dawley and Long Evans rats

**DOI:** 10.64898/2025.12.22.696048

**Authors:** Amber N. Baldwin, Matthew L. Banks

## Abstract

The absence of Food and Drug Administration (FDA)-approved pharmacotherapies for methamphetamine use disorder (MUD) highlights the need for preclinical research to understand both the basic biological mechanisms of methamphetamine reinforcement and evaluate novel MUD pharmacotherapies. Recent studies demonstrated that repeated treatment with the muscarinic M1/M4 receptor agonist xanomeline attenuated cocaine self-administration. Whether these xanomeline treatment effects extend to methamphetamine self-administration remains unknown. The first aim established the economic substitutability between methamphetamine and liquid food (i.e., Ensure®) using a methamphetamine-vs-food choice procedure in male and female Sprague Dawley (SD) and Long Evans (LE) rats. A within-session methamphetamine choice dose-effect function (0.032-0.32 mg/kg/infusion) was determined daily, and food reinforcer magnitude was manipulated weekly by changing the concentration (0, 10, 32, and 100%) of vanilla-flavored Ensure. Additionally, methamphetamine response requirement (i.e., fixed ratio (FR) 1, 5, 25, 125) was manipulated each week while holding the food FR constant. The second aim determined the effectiveness of repeated 5-day xanomeline (3.2-10 mg/kg, SC) to attenuate methamphetamine choice. Both increasing Ensure concentrations and methamphetamine FR values resulted in rightward shifts in the methamphetamine choice dose-effect function in both SD and LE rats. Repeated 5-day xanomeline treatment significantly decreased methamphetamine choice across all doses tested in LE, but not SD, rats. Time course of xanomeline treatment effectiveness revealed effects were greatest during the first 30 min of choice session. These results demonstrate that methamphetamine and food function as economic substitutes and that xanomeline may warrant further consideration as a MUD pharmacotherapy.

## 1.0 Introduction

In wake of the ongoing opioid crisis, illicit methamphetamine use has been simultaneously increasing either by co-use with fentanyl or methamphetamine alone. For example, methamphetamine is now the second leading cause for drug overdoses deaths in the United States(Fischer et al., 2021). Furthermore, methamphetamine was the top reported drug in the September 2023 snapshot report released by The National Forensic Laboratory Information System (System, 2023). Unfortunately, there are currently no available Food and Drug Administration (FDA)-approved pharmacotherapies for Methamphetamine Use Disorder (MUD) highlighting the need for continued preclinical research. Improving translational predictive validity between preclinical animal models to human patients should facilitate translation of our basic understanding of MUD and evaluation of effective candidate MUD medications.

Human drug-taking behavior occurs in an open economy where drug acquisition and use compete with alternative nondrug reinforcers that can range from a diversity of positive options. These alternative nondrug reinforcers include money acquisition through employment, social interaction with friends and family or healthy lifestyle choices(Acuff et al., 2023; Banks and Negus, 2017; Lamb and Ginsburg, 2017). Drug and nondrug reinforcers operate on a continuum of economic commodities(Bickel et al., 1995; Hursh and Silberberg, 2008). Along this continuum, perfect substitutes are at one end and exhibit inverse consumption patterns. Perfect complements are at the opposite end and exhibit identical consumption patterns. Finally, independents fall in the middle and exhibit no correlation in consumption behavior (Kearns, 2025). Nonhuman drug addiction “choice” models capture simplified aspects of this potential economic substitutability when an organism is afforded the opportunity to allocate behavior between two concurrently available reinforcers such as an addictive drug (e.g. methamphetamine) and an alternative nondrug reinforcer (e.g. liquid palatable food) in a controlled operant environment(Townsend et al., 2021). Although addictive drugs such as cocaine, heroin, and fentanyl have been extensively studied in a drug choice context(Kearns et al., 2016; Marcus and Banks, 2025; Townsend et al., 2019; Townsend et al., 2021), there have been a paucity of methamphetamine choice studies in nonhuman primates and rats(Banks and Blough, 2015; Stocco et al., 2025; Venniro et al., 2018). Based on the extant literature with other addictive drugs, we hypothesize that methamphetamine and food will function as economic substitutes. Establishing the sensitivity of methamphetamine-vs-food choice in rats to parametric manipulations should facilitate interpretation of pharmacological treatment efficacy.

One emerging neurotransmitter system that may alter methamphetamine reinforcement is muscarinic acetylcholine receptors. The mesolimbic dopamine “reward” pathway with projections from ventral tegmental area (VTA) dopaminergic neurons terminating in the nucleus accumbens (NAc) has been implicated in reinforcement and connected to executive functioning brain regions such as the prefrontal cortex (PFC) and hippocampus(Holly et al., 2024; Russo and Nestler, 2013). M1 G_q_-protein coupled receptors are expressed on medium spiny neurons in the NAc, and broadly expressed broadly throughout the PFC(Hersch et al., 1994). In addition, M4 Gi-protein coupled receptors are expressed throughout the caudate-putamen and NAc on medium spiny neurons and signal through inhibitory cascades(Weiner et al., 1990). In preclinical behavioral studies, repeated treatment with the M1/M4-preferring agonist xanomeline significantly decreased cocaine self-administration in a cocaine-vs-food choice procedure(Marsh et al., 2025; Thomsen et al., 2014). Furthermore, the M1 bitopic agonist VU0364572 decreased cocaine-vs-food choice in male rats(Weikop et al., 2020). Overall, these results suggest the potential of muscarinic ligands to modulate psychostimulant self-administration towards the development of candidate pharmacotherapies.

The aim of the present study was to determine the effects of three nonpharmacological manipulations and one pharmacological manipulation on methamphetamine-vs.-food choice in male and female rats. Alternative nondrug reinforcer magnitude and response requirement manipulations were determined because previous cocaine (Negus, 2003; Thomsen et al., 2013) and opioid (St. Onge et al., 2022; Townsend et al., 2021) choice studies in rats demonstrate that these experimental manipulations should alter drug choice. These nonpharmacological manipulations also establish the boundary conditions of the procedure and facilitate interpretation of pharmacological manipulations(Banks and Negus, 2017). The third nonpharmacological manipulation was to compare alternative nondrug reinforcer and response requirement manipulations between two common rat strains used in drug self-administration studies. There is a small literature suggesting potential strain differences between Sprague Dawley and Long Evans rats that might influence methamphetamine choice(Swerdlow et al., 2006). Lastly, we determined the effects of repeated 5-day xanomeline treatment on methamphetamine-vs-food choice in both Sprague Dawley and Long Evans rats under a one-hour behavioral session due to the short duration of action of xanomeline(Bymaster et al., 1997; Mirza et al., 2003). Xanomeline treatment-induced decreases in methamphetamine choice across both rat strains would provide robust evidence supporting the continued consideration of xanomeline, clinically available with the peripheral muscarinic antagonist trospium (Cobenfy ©) as a candidate MUD pharmacotherapy.

## 2.0 Methods

### 2.1 Subjects

Adult male and female Sprague Dawley (n = 6-8M/5-6F) and Long Evans (n=6-7M/6F) rats weighing 250-350 grams upon delivery (Envigo, Frederick, MD) served as research subjects. Subjects were fed standard laboratory chow (Teklad Rat Diet, Envigo) ad-libitum with continuous water access in their home cage. Animals were singly housed in an AAALAC-accredited temperature- and humidity-controlled vivarium that contained a 12-h light/dark cycle (Lights off at 6:00PM). All rats were surgically implanted with an indwelling jugular catheter and vascular access port using aseptic techniques as previously published(Townsend et al., 2021). Behavioral sessions occurred 5 days a week from approximately 1 pm – 4 pm. Animal maintenance and research were conducted in accordance with the 2011 Guidelines of the National Institutes of Health Committee on Laboratory Animal Resources. Both enrichment and research protocols were approved by the Virginia Commonwealth University Institutional Animal Care and Use Committee.

### 2.2 Drugs

(+)-Methamphetamine HCl was provided through the NIDA Drug Supply Program (RTI International, RTP, NC). Methamphetamine was dissolved in sterile water and passed through a 0.22 µm sterile filter (Millex GV, Millipore Sigma, Burlington, MA) before intravenous (IV) administration. Xanomeline oxalate was purchased from a commercial vendor (Tocris, Minneapolis, MN or MedChemExpress, Monmouth Junction, NJ) and was dissolved in bacteriostatic saline. All drug doses are expressed as the salt forms listed above.

### 2.3 Apparatus and Catheter maintenance

Modular operant chambers located in sound-attenuating cubicles (Med Associates, St. Albans, VT, USA) were equipped with two retractable levers, a set of three LED lights (red, yellow, green) mounted above each lever, and a retractable “dipper” cup (0.1 ml) located between the levers for presenting diluted Ensure® (10%, 32% and 100% v/v vanilla-flavored Ensure® in tap water; Abbott Laboratories, Chicago, IL, USA). In addition, a syringe pump (PHM-100, Med Associates) located inside the sound-attenuating cubicle delivered IV methamphetamine infusions using custom Med-PC programming as previously published(Townsend et al., 2021). After each behavioral session, catheters were flush with a 24-h gentamicin (0.04 mg/0.1 ml) lock solution. Catheter patency was verified at the end of each experiment by confirmation of instantaneous muscle tone loss following IV methohexital (0.5 mg/0.1 ml) administration.

### 2.4 Within-session dose methamphetamine-vs.-food choice procedure

The terminal IV methamphetamine-vs.-food self-administration session was approximately 2h in duration and consisted of a concurrent fixed-ratio (FR)5:FR5 schedule of IV methamphetamine and food availability. There were five successive response components and a different ascending IV methamphetamine dose (0, 0.01, 0.032, 0.1, and 0.32 mg/kg/infusion per weekly body weight of subject) was available as the alternative to liquid food in each successive component. Methamphetamine dose was manipulated by changing the infusion duration(Townsend et al., 2021). Each response component was 20-min with a 5-min timeout period separating each response component. Within each response component, subjects could complete up to 10 response requirement completions or “choices” on either the liquid food or methamphetamine-associated lever. Methamphetamine-vs-food choice behavior was considered stable when the smallest unit methamphetamine dose that maintained greater than 80 percent methamphetamine choice did not vary by more than 0.5 log units across three sessions.

Once methamphetamine-vs-food choice stabilized for an individual subject, parametric manipulations were initiated starting the Monday of the following week and ending on Friday. Two different experiments were conducted. First, the magnitude of the liquid food reinforcer was manipulated by changing the concentration of vanilla-flavored Ensure® using tap water (0, 10, 32, and 100%) each week. Second, the response requirement for methamphetamine was manipulated, with each FR value (1, 5, 25, and 125) determined each week, while holding the response requirement for food at FR5. These experimental manipulations were determined in a counterbalanced fashion both within an experiment and between the two experiments.

### 2.5 Single dose methamphetamine-vs.-food choice procedure

A one-hour free operant methamphetamine-vs.-food choice procedure was used to evaluate repeated 5-day xanomeline treatment effects. The one-hour IV methamphetamine-vs.-food self-administration session was a concurrent FR5:FR5 schedule of a single methamphetamine dose (0.1 or 0.32 mg/kg/infusion) or liquid food (100% for Sprague Dawley and 32% for Long Evans rats) availability. These Ensure concentrations were based on the within-session choice results above. Xanomeline (3.2, 5.6, and 10 mg/kg, SC) and vehicle were administered as a five-minute pretreatment Monday through Friday for five consecutive days. Baseline weeks with no treatments separated each treatment week and xanomeline doses were counterbalanced between rats.

### 2.6 Data Analysis

The primary dependent measure was percent methamphetamine choice defined as the number of methamphetamine choices per component ÷ the total number of choices per component × 100. These data were plotted as a function of methamphetamine dose and independent variable manipulation (i.e. ensure concentration or methamphetamine FR) using GraphPad Prism (Prism 10, GraphPad, La Jolla, CA, USA). Additional dependent measures included session total, methamphetamine, and food choices. Furthermore, percent methamphetamine choice data were transformed by plotting unit price (FR÷ dose) on the abscissa and fit using the Zero-Bounded Exponential Model of Demand (Hursh and Silberberg, 2008) via a GraphPad Prism template (available from the Institutes for Behavior Resources, http://www.ibrinc.org). This transformation facilitated calculation of both Q_0_ defined as consumption at unconstrained price and essential value (EV) defined as reinforcer strength similar to previous studies(Hursh and Silberberg, 2008; St. Onge et al., 2024; Townsend et al., 2019). Statistical comparisons utilized a mixed-effects analysis, and the Geisser-Greenhouse correction was applied as appropriate for within-subject comparisons. Statistical significance was established *a priori* as p<0.05.

## 3.0 Results

### 3.1 Effects of manipulating Ensure concentration on methamphetamine choice

Figure 1 shows the effects of manipulating the reinforcer magnitude on methamphetamine-vs.-food choice in Sprague Dawley and Long Evans male and female rats. Methamphetamine maintained a dose-dependent increase in percent methamphetamine choice in both strains. In Sprague-Dawley rats (Figure 1A), decreasing the Ensure concentration from 100 percent to zero percent (i.e., water) resulted in a concentration-dependent increase in percent methamphetamine choice (methamphetamine dose: F_2.11, 23.20_ = 30.09, p < 0.0001; Ensure concentration: F_2.84, 31.22_ = 0.95, p < 0.0001 ; interaction: F_3.06, 22.09_ = 0.34, p = 0.02). Figure 1B shows the effects of manipulating reinforcer magnitude on methamphetamine-vs.-food choice in Long Evans rats. Similar to Sprague Dawley rats, decreasing the Ensure concentration from 100 percent to zero percent resulted in a concentration-dependent increase in percent methamphetamine choice (methamphetamine dose: F_2.15, 23.64_ = 0.72, p < 0.0001; Ensure concentration: F_2.49, 27.43_ = 0.83, p = 0.0002, interaction: F_3.89, 31.57_ = 0.43, p = 0.024).

**Figure 1.**
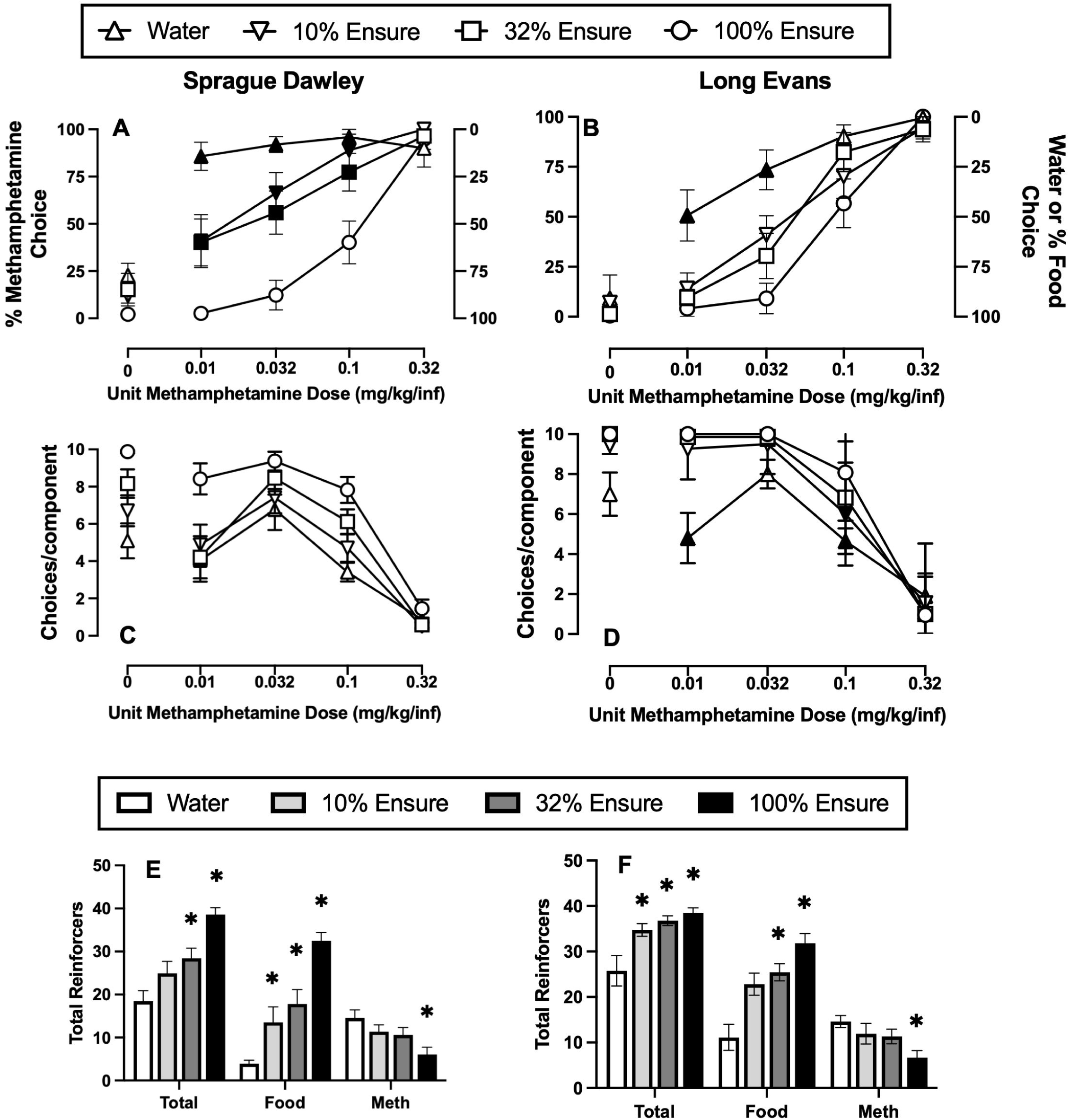
Effect of liquid food concentration manipulations on methamphetamine choice in Sprague Dawley (SD) and Long Evans (LE) rats. Panels A (SD) and B (LE) show percent methamphetamine choice as a function of unit methamphetamine dose in milligrams per kilogram per infusion. Panels C (SD) and D (LE) show the number of choices completed per component. Panels E (SD) and F (LE) show session total, food, and methamphetamine reinforcers. All data represent the mean ± SEM for SD (n = 6M/6F) and LE (n = 6-7M/6F) rats. Filled symbols and asterisks denote a significant difference between Ensure concentration and water.

Figure 1C-D shows the number of choices completed per each component of the methamphetamine choice procedure in Sprague Dawley (1C) and Long Evans (1D) rats. Across both strains, the number of choices completed per component decreased as a function of increasing methamphetamine doses available. Manipulating the Ensure concentration did not significantly alter the choices completed per component in Sprague Dawley rats (methamphetamine dose: F_1.79, 19.68_ = 53.10, p < 0.0001; Ensure concentration: F_2.68, 29.45_ = 15.82, p < 0.0001; interaction: F_4.17, 44.04_ = 0.46, p = 0.088). In Long Evans rats, when water was the alternative to methamphetamine infusions, choices completed per component at 0.01 mg/kg/infusion methamphetamine dose was significantly different from 100 percent Ensure (methamphetamine dose: F_2.15, 23.63_ = 86.11, p < 0.0001; Ensure concentration: F_2.49, 27.43_ = 10.38, p = 0.0002; interaction: F_3.11, 28.32_ = 0.35, p = 0.006). In addition, water and 10 percent Ensure were significantly different from 100 percent Ensure at the 0.1 mg/kg/infusion methamphetamine dose.

Figure 1D and 1E shows session total, methamphetamine, and food reinforcers earned during the 2-h behavioral session in Sprague Dawley (1D) and Long Evans (1E) rats, respectively. As the Ensure concentration increased, the number of food reinforcers increased and conversely the number of methamphetamine reinforcers decreased without altering session total reinforcers in Sprague Dawley rats (Ensure concentration: F_1.18, 12.99_ = 32.42, p < 0.0001; reinforcer type: F_2.13, 23.41_ = 26.05, p < 0.0001; interaction: F_1.98, 18.77_ = 24.12, p < 0.0001). In Long Evans rats, decreasing Ensure concentrations significantly decreasing session food reinforcers and correspondingly increasing session methamphetamine reinforcers (Ensure concentration: F_2.15, 23.64_ = 86.11, p < 0.0001; reinforcer type: F_2.49, 27.43_ = 26.05, p = 0.0002; interaction: F_2.38,16.63)_ = 11.34, p = 0.0005). In addition, water availability significantly reduced total reinforcers compared to all Ensure concentrations.

When comparing Ensure concentration manipulations between Sprague Dawley and Long Evans, there were no statistical differences between rat strains for any Ensure concentration.

### 3.2 Effects of methamphetamine FR manipulations on methamphetamine choice

Figure 2 shows the effects of selectively manipulating the methamphetamine FR value on methamphetamine-vs.-food choice in Sprague Dawley (Figure 2 A, C) and Long Evans Rats (Figure 2 B, D). In both strains, when the FR values were equal for both methamphetamine and food at FR5, methamphetamine maintained a dose-dependent increase in percent methamphetamine choice. In Sprague Dawley rats (Figure 2A), increasing the methamphetamine FR value significantly decreased 0.032 mg/kg/infusion methamphetamine choice at FR5, 25, and 125 compared to FR1, 0.1 mg/kg/infusion methamphetamine choice at FR25 and 125, and 0.32 mg/kg/infusion methamphetamine choice at FR125 (dose: F_2.24, 26.85_ = 23.92, p < 0.0001; FR: F_2.00, 23.95_ = 18.57, p < 0.0001; interaction: F_3.83, 29.88_ = 0.43, p = 0.001). There was no significant effect of manipulating the methamphetamine FR value on choices completed per component (dose: F_2.25, 27.00_ = 89.71, p < 0.0001; FR: F_2.14, 25.62_ = 0.54, p = 0.599; interaction: F_4.43, 41.34_ = 2.41, p = 0.059) (Figure 2C). In Long Evans rats (Figure 2B), 0.032 and 0.1 mg/kg/infusion methamphetamine choice was significantly decreased at FR 25 and 125, and 0.32 mg/kg/infusion methamphetamine choice was significantly decreased at FR125 (dose: F_1.86, 22.33_ = 41.19, p < 0.0001; FR: F_2.49, 29.89_ = 31.42, p < 0.0001; interaction: F_2.91, 20.01_ =0.32, p=0.006). Although there was a significant methamphetamine dose × FR value interaction on choices completed per component (dose: F_2.41, 28.95_ = 80.86, p < 0.0001; FR: F_2.07, 24.78_ = 0.86, p = 0.437; interaction: F_3.93, 36.64_ =0.44, p=0.045) in Long Evans rats (Figure 2D), no significance differences were detected post-hoc following multiple comparison corrections.

**Figure 2.**
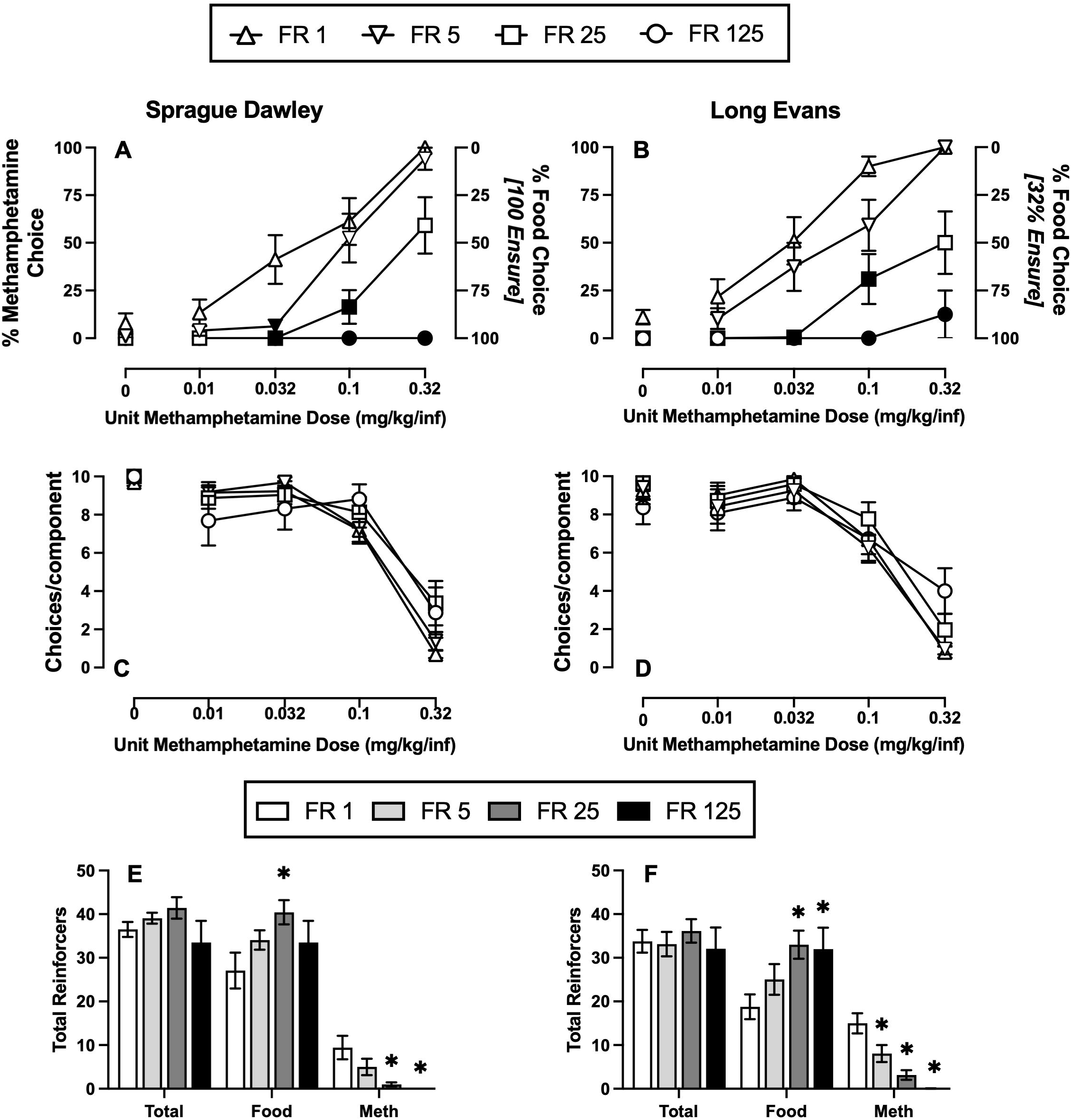
Effect of methamphetamine FR manipulations on methamphetamine choice in Sprague Dawley (SD) and Long Evans (LE) rats. Panels A (SD) and B (LE) show percent methamphetamine choice as a function of unit methamphetamine dose in milligrams per kilogram per infusion. Panels C (SD) and D (LE) show the number of choices completed per component. Panels E (SD) and F (LE) show effects of methamphetamine FR manipulations on session total, food, and methamphetamine reinforcers. All data represent the mean ± SEM of SD (6-8M/ 2-6F) and LE (4-5M/ 6-7F) rats. Filled symbols and asterisks denote a significant difference compared to FR 1.

Figure 2E-F shows session total, food, and methamphetamine reinforcers in Sprague Dawley and Long Evans, respectively. In Sprague Dawley rats, session food reinforcers were significantly increased when the methamphetamine FR was 25 compared to FR1 and session methamphetamine reinforcers were significantly decreased at methamphetamine FR values of 25 and 125 (FR: F_1.25, 7.50_ = 119.3, p < 0.0001; reinforcer type: F_1.82, 10.90_ = 1.85, p = 0.204; interaction: F_2.79, 15.33_ = 0.48, p=0.017). In Long Evans, session food reinforcers were significantly increased when the methamphetamine FR was 25 and 125 compared to FR1 and session methamphetamine reinforcers were decreased at methamphetamine FR values of 5, 25, and 125 (FR: F_1.55, 15.48_ = 48.64, p < 0.0001; reinforcer type: F_1.42, 14.22_ = 0.60, p = 0.506; interaction: F_2.48,18.60=_ 25.42, p < 0.0001).

Figure 3 compares methamphetamine choice demand curves between Sprague Dawley and Long Evans rats by transforming the data shown in Figure 2 using the Zero-Bounded Exponential Model(Hursh and Silberberg, 2008). There was no significant effect of rat strain on either methamphetamine consumption at unconstrained price (mean Q_0_, 95% confidence interval; Sprague Dawley: 199, -9.4 to 408; Long Evans: 332: 9.5 to 654) or methamphetamine reinforcer strength (mean essential value (EV), 95% confidence interval; Sprague Dawley: 231, 41.6 to 421; Long Evans: 634, 142 to 1126).

**Figure 3.**
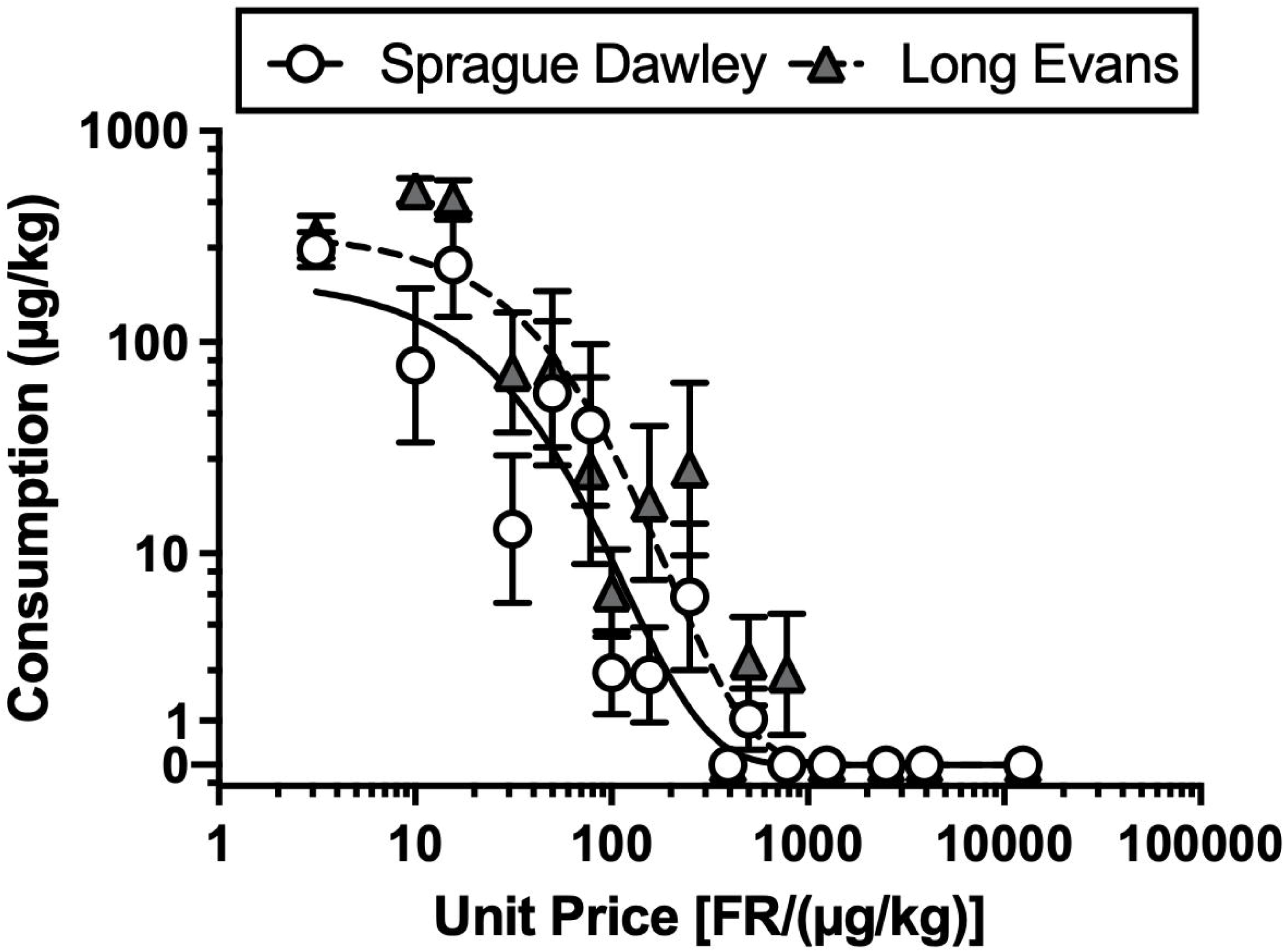
Methamphetamine-vs-food choice demand curve in Sprague Dawley (SD) and Long Evans (LE) Rats. Data are replotted from Figure 2A and 2B and represent the mean ± SEM for both SD (6-8M/2-5F) and LE (5-7M/2-5F) rats.

### 3.3 Methamphetamine-vs.-food choice under free operant 1-h procedure

Because of the short duration of action of xanomeline, a one-hour methamphetamine-vs.-food choice procedure was developed, and the methamphetamine dose-effect function is shown in Figure 4 for Sprague Dawley (A, C) and Long Evans (B, D) rats. Percent methamphetamine choice increased as a function of methamphetamine dose in both Sprague Dawley (4A) and Long Evans (4B) rats. Methamphetamine doses of 0.032, 0.1, and 0.32 mg/kg/infusion maintained significantly greater methamphetamine choice than saline in both Sprague Dawley (methamphetamine dose: F_4,33_ = 4.11, p < 0.0001) and Long Evans rats (methamphetamine dose: F_4,23_ = 1.330, p = 0.0003). As with the within-session dose-effect choice procedure, there were no significant differences between rat strains (methamphetamine dose: F_4, 55_: 24.86, p < 0.0001; strain: F_1, 55_: 1.15, p = 0.288; interaction: F_4, 55_: 0.58, p = 0.680). Figure 4C and 4D shows the number of methamphetamine and food reinforcers earned in Sprague Dawley and Long Evans rats, respectively. As the methamphetamine dose increased, methamphetamine infusions significantly increased at methamphetamine doses of 0.01 and 0.032 mg/kg/infusion compared to saline and food reinforcers significantly decreased at all methamphetamine doses examined (reinforcers: F_1.00,66.00_ = 1.18, p = 0.282; dose: F_1.43, 23.62_ = 13.03, p = 0.0005; interaction: F_1.85, 30.56_ = 16.17, p < 0.0001). In Long Evans rats, food reinforcers were significantly decreased at all methamphetamine doses compared to saline (reinforcers: F_1.00,6.00_ = 3.61, p = 0.106; dose: F_1.36, 8.17_ = 10.64, p = 0.008; interaction: F_1.62, 6.46_ = 8.23, p = 0.019).

**Figure 4.**
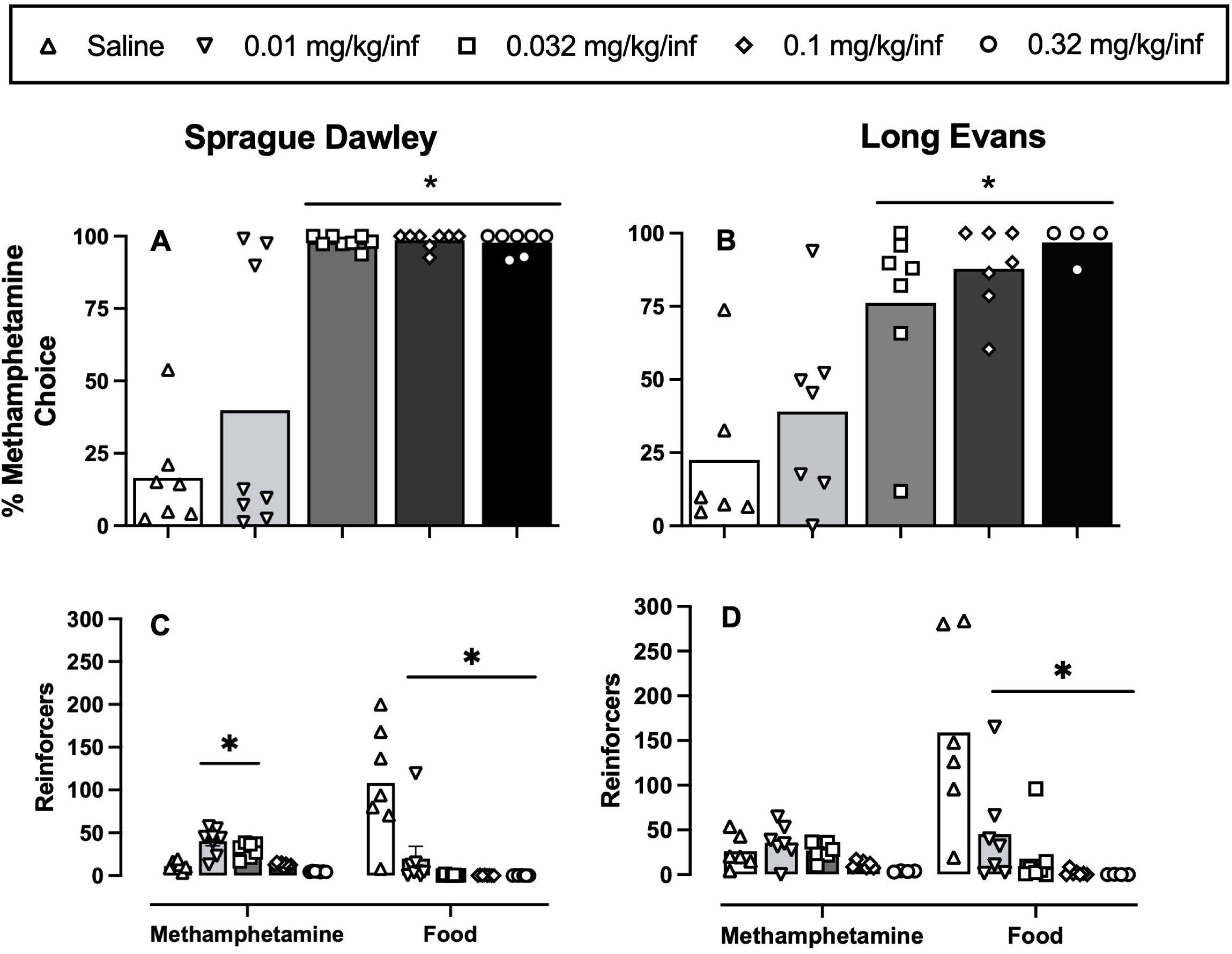
Methamphetamine-vs-food choice as a function of methamphetamine dose in a 1-h free operant procedure in Sprague Dawley (SD) and Long Evans (LE) rats. Panels A (SD) and B (LE) show percent methamphetamine choice as a function of unit methamphetamine dose in milligrams per kilogram per infusion. Panels C (SD) and D (LE) show session methamphetamine and food reinforcers. All data represent the mean for SD (6-8M/2-5F) and LE (5-7M/2-5F) rats with symbols denoting individual subject data. Dash above bars denote statistical significance (p<0.05) compared to saline.

#### Xanomeline treatment effects on 0.1 mg/kg/infusion methamphetamine choice

Figure 5 A-D shows repeated 5-day xanomeline (3.2, 5.6, and 10 mg/kg/day, SC) treatment effects on 0.1 mg/kg/infusion methamphetamine choice and session methamphetamine and food reinforcers in Sprague Dawley (A, C) and Long Evans (B, D) rats. No xanomeline treatment dose significantly altered methamphetamine choice or session methamphetamine reinforcers in Sprague Dawley rats. In contrast to Sprague Dawley rats, all three xanomeline doses (3.2, 5.6, and 10 mg/kg/day) significantly attenuated methamphetamine choice (xanomeline dose: F_1.693,9.591_ = 13.03, p = 0.0024). Furthermore, all xanomeline doses significantly increased food reinforcers without significantly altering methamphetamine reinforcers (reinforcer type: F_1.00, 6.00_ = 3.61, p = 0.106; xanomeline dose: F_1.36, 8.17_ = 10.64, p = 0.008; interaction: F_1.62,6.46_ = 8.23, p = 0.019).

**Figure 5.**
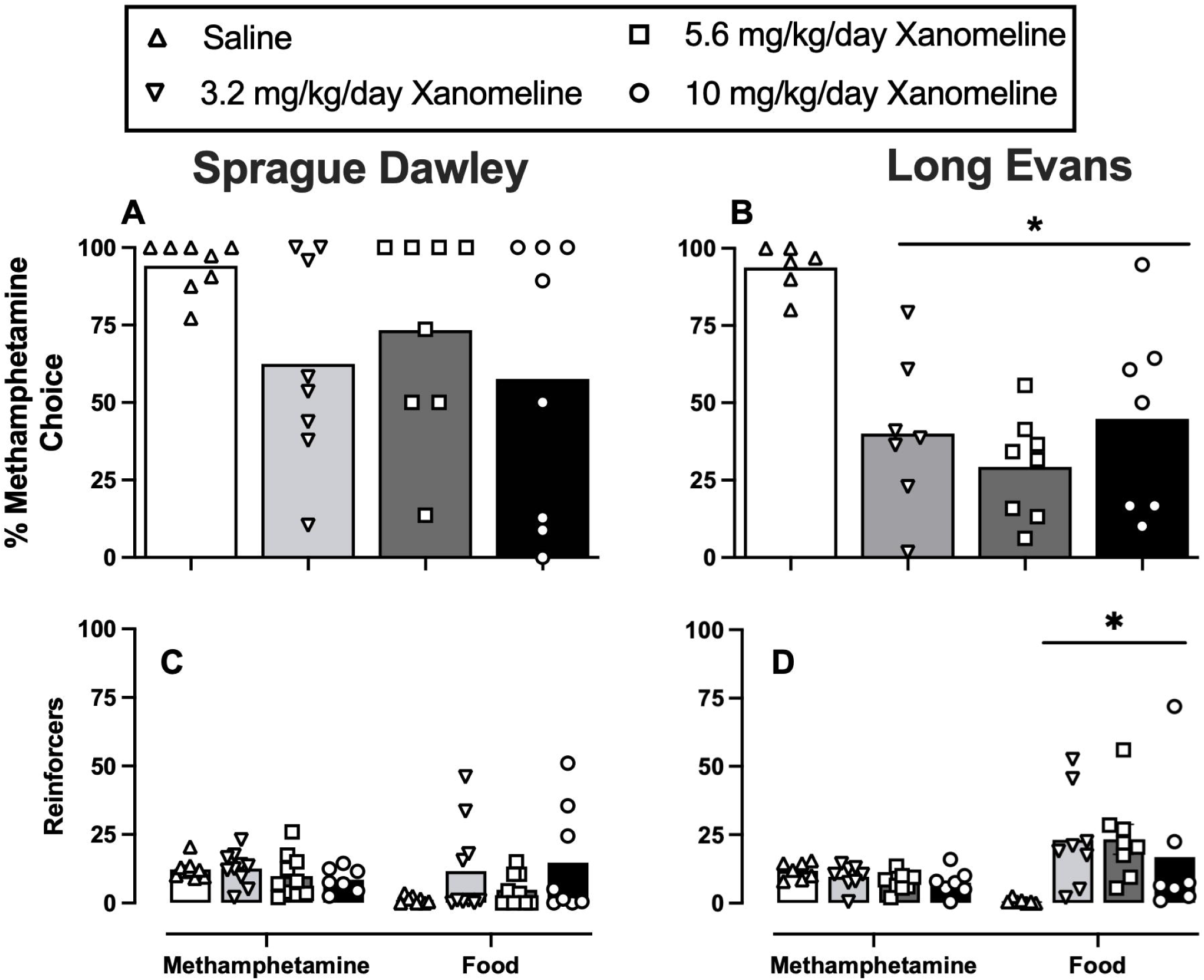
Repeated 5-day xanomeline treatment effects on 0.1 mg/kg/inf methamphetamine choice in Sprague Dawley (SD) and Long Evans LE) rats. Panels A (SD) and B (LE) show percent methamphetamine choice as a function of xanomeline treatment dose (0, 3.2-10 mg/kg/day, SC). Panels C (SD) and D (LE) show session methamphetamine and food reinforcers. All data represent the mean for SD (6-8M/2-5F) and LE (5-7M/2-5F) rats with symbols denoting individual subject data. Dash above bar denotes statistical significance (p<0.05) compared to saline.

### 3.4 Time course of xanomeline treatment effects

Figures 6 and 7 show the time course of xanomeline treatment effects across 15-min time bins in Sprague Dawley and Long Evans rats, respectively. In Sprague Dawley rats, there was no significant bin × day interaction for either food or methamphetamine during 3.2 mg/kg/day xanomeline treatment (food reinforcers per bin: F_1.19,10.72_ = 5.73, p = 0.032; day: F_2.40, 21.63_ = 2.00, p = 0.152; interaction: F_4.38, 39.43_ = 1.69, p = 0.166; methamphetamine reinforcers per bin: F_1.27,11.47_ = 10.06, p = 0.006; day: F_3.06, 27.49_ = 1.00, p = 0.411; interaction: F_4.68, 42.12_ = 1.87, p = 0.124). For 5.6 mg/kg/day xanomeline treatment, there was no significant food reinforcers per bin × day interaction; however, xanomeline significantly decreased methamphetamine reinforcers in the first 15-min bin at day 3 compared to day 0 (methamphetamine reinforcers per bin: F_1.58, 14.18_ = 2.11, p = 0.163; day: F _3.36, 30.25_ = 1.21, p = 0.327; interaction: F_4.43, 39.88_ = 3.51, p = 0.023). For 10 mg/kg/day xanomeline, there was no significant bin × day interaction for either food or methamphetamine reinforcers in Sprague Dawley. For Long Evans, there was no significant food reinforcers per bin × day interaction (food reinforcers per bin: F_1.15, 8.04_ = 14.76, p = 0.004; day: F _2.22, 15.54_ = 5.27, p = 0.016; interaction: F_3.00, 21.00_ = 2.64, p = 0.076); however, there was a significant decrease in methamphetamine reinforcers in the first 15-min bin at days 3, 4, and 5 compared to day 0 (methamphetamine reinforcers per bin: F _2.35, 16.47_ = 3.79, p = 0.039; day: F _2.33, 16.34_ = 0.986, p = 0.406; interaction: F_3.63, 25.44_ = 3.43, p = 0.026). 5.6 mg/kg xanomeline significantly increased food reinforcers in the first 15-min bin at day 3 compared to day 0 (food reinforcers per bin: F _1.11, 7.76_ = 6.55, p = 0.033; day: F _2.58, 18.08_ = 9.52, p = 0.001; interaction: F_2.58, 18.04_ = 3.84, p = 0.032) and significantly decreased methamphetamine reinforcers in the first 15-min bin at days 1-5 compared to day 0 (methamphetamine reinforcers per bin: F _1.86, 13.02_ = 1.38, p = 0.285; day: F _2.79, 19.52_ = 2.54, p = 0.089; interaction: F_2.58, 18.04_ = 3.84, p = 0.032). Although 10 mg/kg xanomeline did not significantly increase food reinforcers (food reinforcers per bin: F _1.10, 6.58_ = 2.00, p = 0.205; day: F _1.43, 8.56_ = 2.45, p = 0.151; interaction: F_1.23, 7.79_ = 1.80, p = 0.225), it significantly decreased methamphetamine reinforcers in the first 15-min bin at days 1-5 compared to day 0 (methamphetamine reinforcers per bin: F _2.12, 14.84_ = 2.26, p = 0.138; day: F _1.96, 13.75_ = 4.87, p = 0.026; interaction: F_3.32, 21.47_ = 8.38, p = 0.001).

**Figure 6.**
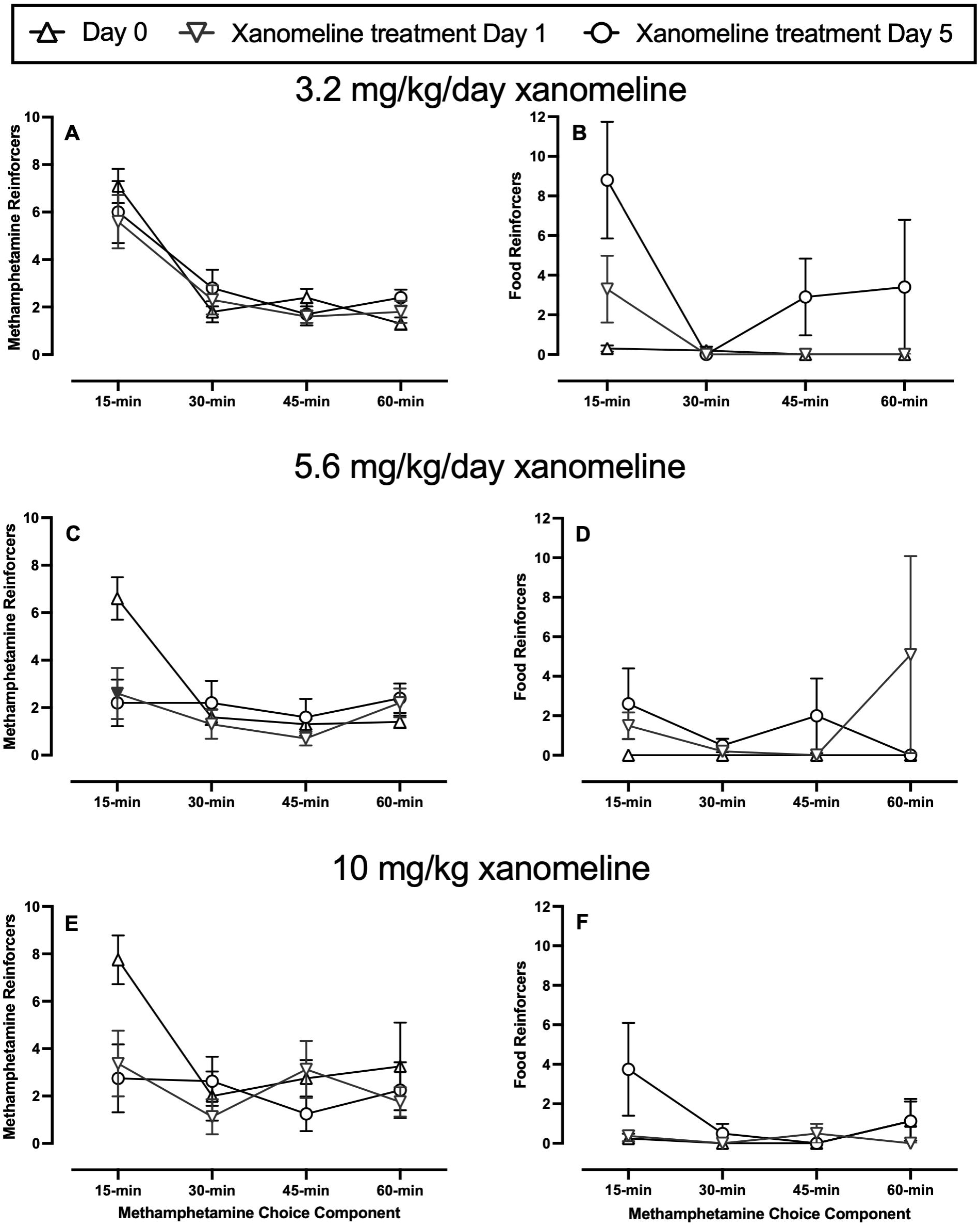
Time course of xanomeline treatment effects both across the five consecutive treatment days and within the 1-h methamphetamine choice procedure in Sprague Dawley rats. Panels A, C, E show methamphetamine reinforcers and panels B, D, E show food reinforcers during 3.2 mg/kg/day (A, B), 5.6 mg/kg/day (C, D), and 10 mg/kg/day (E, F) xanomeline treatment, respectively. Filled symbols denote a statistical significance compared to Day 0 (pre-xanomeline treatment).

**Figure 7.**
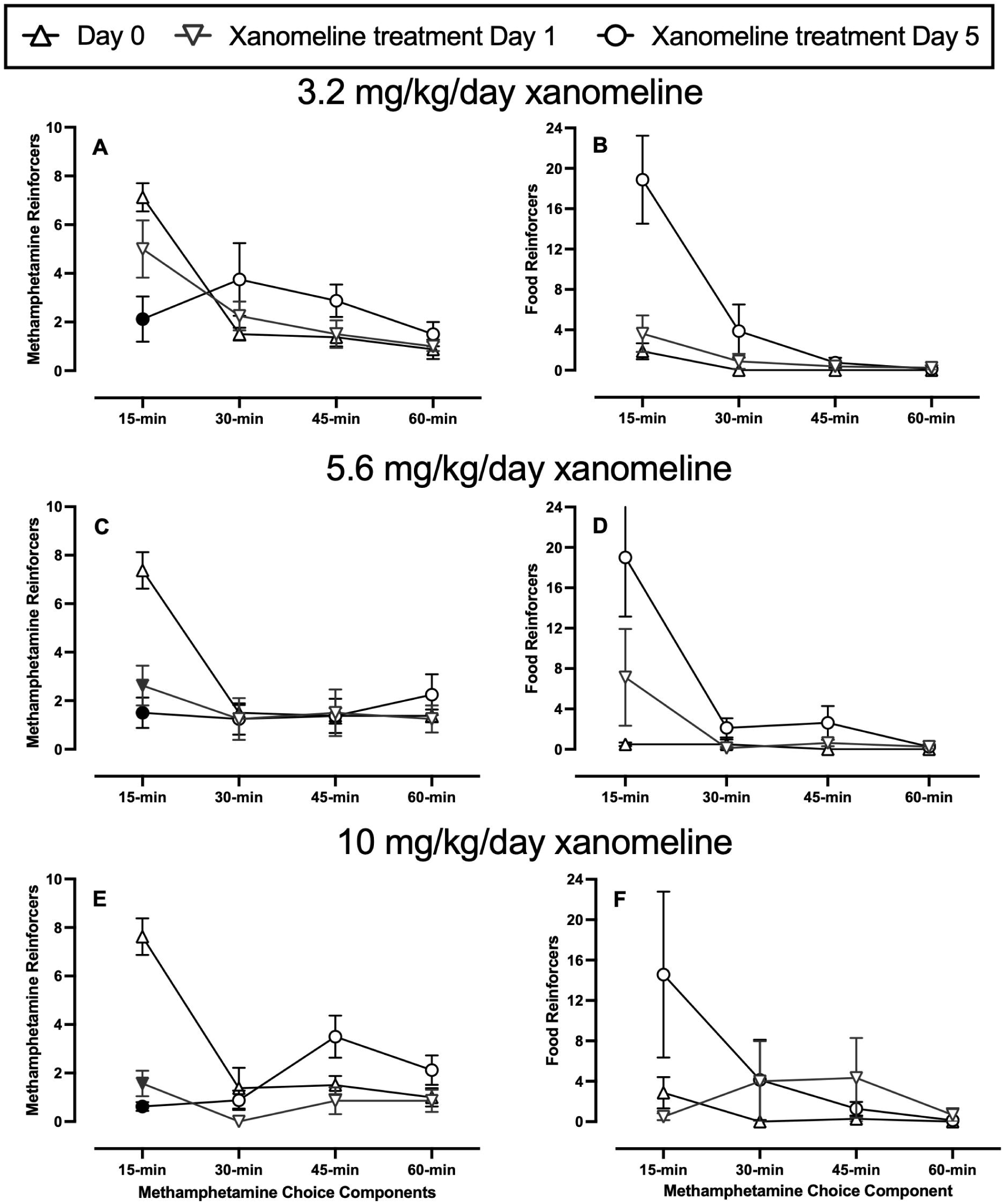
Time course of xanomeline treatment effects both across the five consecutive treatment days and within the 1-h methamphetamine choice procedure in Long Evans rats. Panels A, C, E show methamphetamine reinforcers earned and panels B, D, E show food reinforcers earned for 3.2 mg/kg/day (A, B), 5.6 mg/kg/day (C, D), and 10 mg/kg/day (E, F) xanomeline treatment, respectively. Filled symbols denote a statistical significance compared to Day 0 (pre-xanomeline treatment).

Analysis of xanomeline treatment effects for the first 30 min of the methamphetamine choice session also failed to detect a significant xanomeline effect on methamphetamine choice in Sprague Dawley rats (Figure 8A). However, both 5.6 and 10 mg/kg/day xanomeline significantly decreased methamphetamine reinforcers without significantly increasing food reinforcers during the first 30-min (Figure 8C; reinforcer type: F_1.00, 9.00_ = 0.99, p = 0.346; xanomeline dose: F_2.43, 21.86_ = 2.63, p = 0.085; interaction: F_2.14, 14.97_ = 4.76, p = 0.023). In Long Evans rats (Figure 8B), all three xanomeline doses significantly decreased percent methamphetamine choice (xanomeline dose: F_3.00,26.00_ = 18.10, p < 0.0001). Additionally, 5.6 and 10 mg/kg/day xanomeline significantly decreased methamphetamine reinforcers (Figure 8D; reinforcer type: F_1.00, 7.00_ = 3.28, p = 0.113; xanomeline dose: F_1.54, 10.74_ = 2.52, p = 0.134; interaction: F_1.99, 11.25_ = 6.50, p = 0.013).

**Figure 8.**
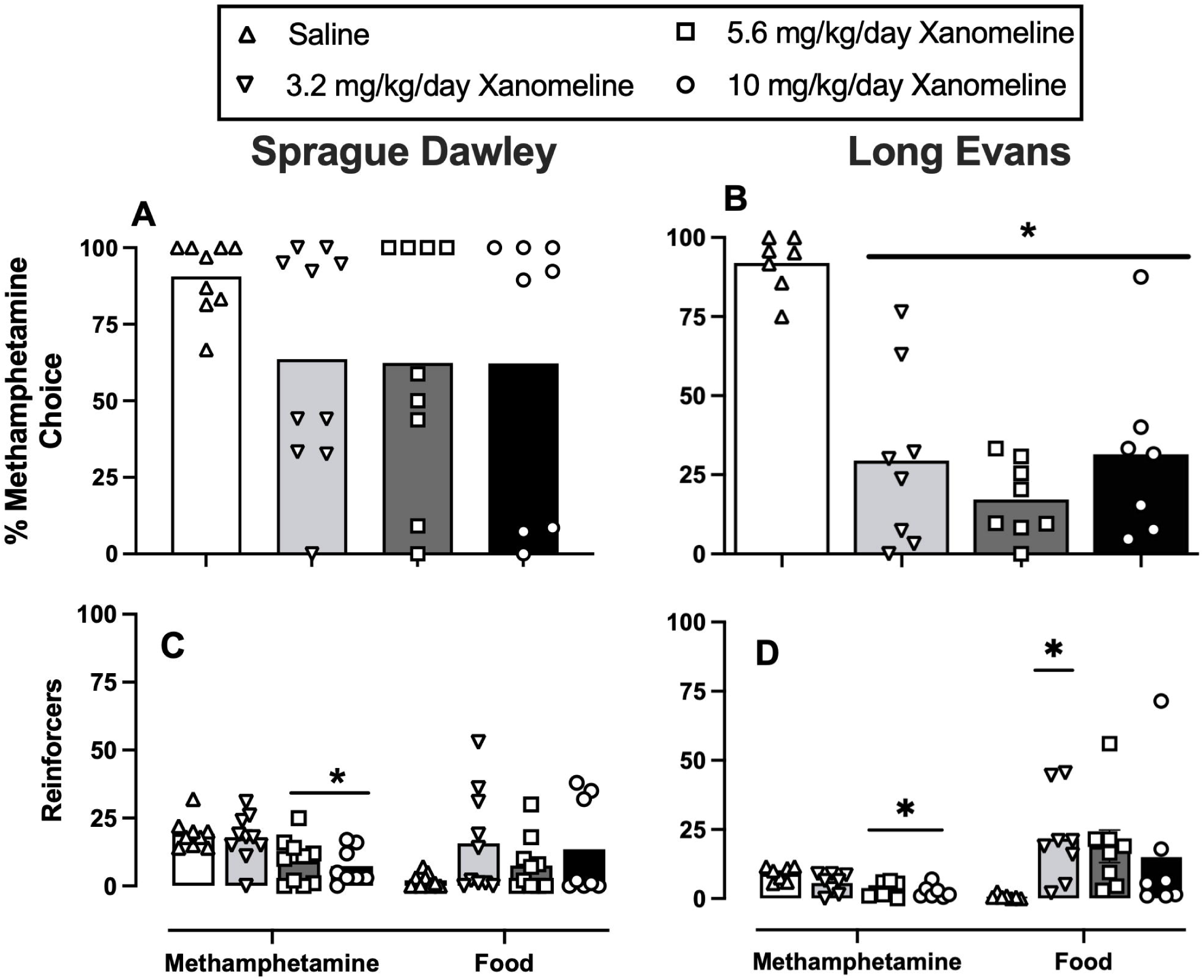
Effects of repeated 5-day xanomeline (3.2-10 mg/kg/day) or saline on 0.1 mg/kg/infusion methamphetamine versus liquid food choice in the first 30-min in Sprague Dawley (SD) and Long Evans (LE) rats. Panels A (SD) and B (LE) show percent methamphetamine choice as a function of xanomeline treatment dose. Panels C (SD) and D (LE) show session methamphetamine and food reinforcers. All bars represent the mean for Sprague Dawley (6-8M/2-5F) and Long Evans (5-7M/2-5F) rats and symbols denote individual subject data. Dash above bar represent statistical significance compared to saline.

### 3.5 Xanomeline treatment effects on 0.32 mg/kg/infusion methamphetamine choice

Figure 9 A-B shows the effects of repeated 5.6 mg/kg/day xanomeline treatment in a mixed cohort of Sprague Dawley and Long Evans rats (n = 7 Sprague Dawley / 3 Long Evans) when a larger 0.32 mg/kg/infusion methamphetamine dose was available as the alternative to liquid food. 5.6 mg/kg/day xanomeline did not significantly attenuate methamphetamine choice (t_9_ = 1.93, p = 0.086; Figure 9A). Neither methamphetamine nor food reinforcers were significantly altered by xanomeline treatment (Figure 9B).

**Figure 9.**
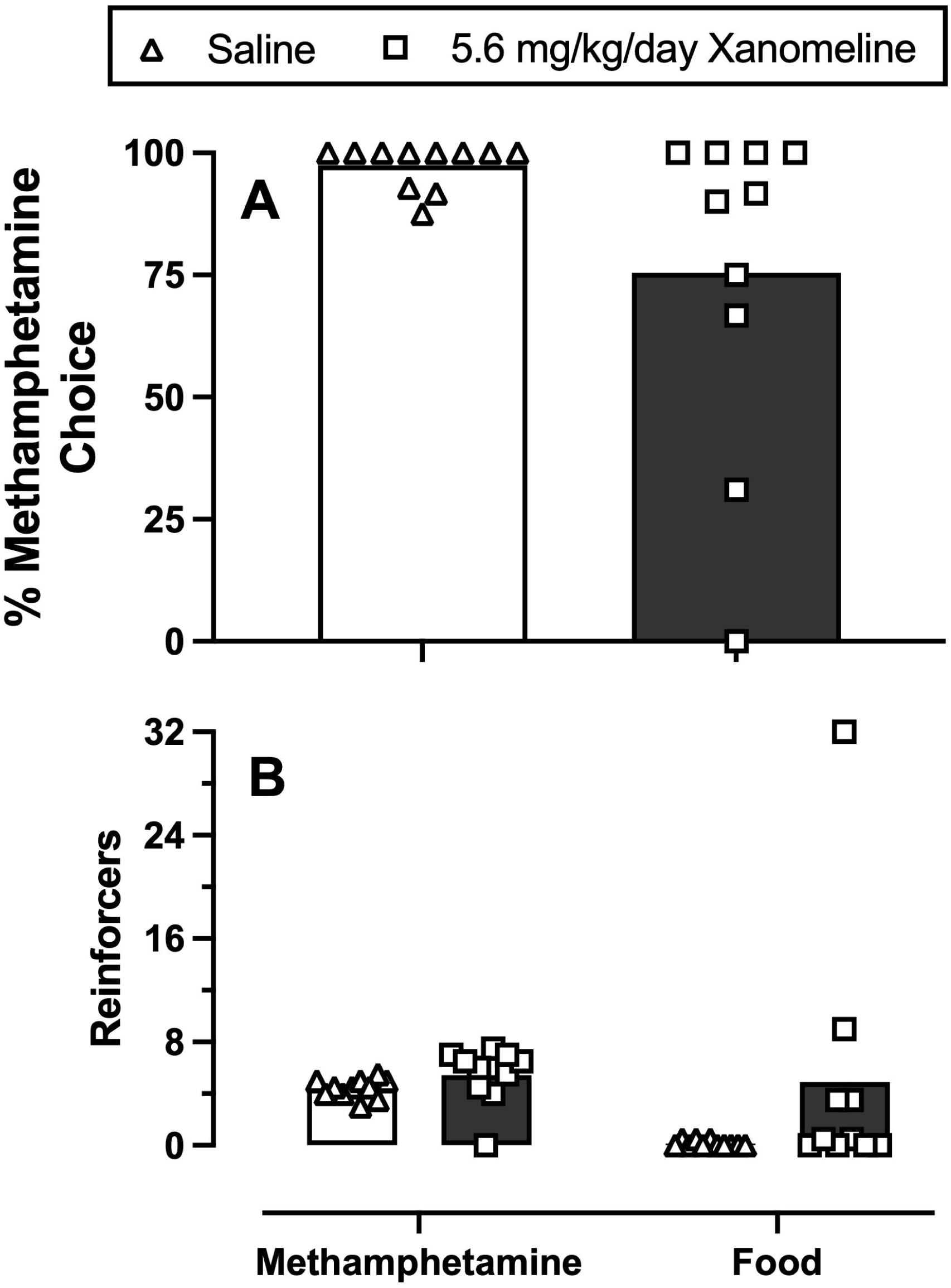
Repeated five-day 5.6 mg/kg xanomeline treatment on 0.32 mg/kg/inf methamphetamine choice. Panel A represents percent methamphetamine choice in a mixed cohort for Sprague Dawley and Long Evans rats (n = 7 SD/3 LE). Panel B shows session methamphetamine and food reinforcers. All bars represent mean data and symbols denote individual subject data.

## 4.0 Discussion

### 4.1 Implications of methamphetamine and food as economic substitutes

The present study explored the behavioral economic and pharmacological determinants of methamphetamine choice in Sprague Dawley and Long Evans male and female rats. There were three main findings from the behavioral economic studies. First, methamphetamine maintained a dose-dependent increase in choice in both Sprague Dawley and Long Evans rats under both the within-session dose-effect and single dose choice procedure consistent with previously published rat (Stocco et al., 2025) and nonhuman primate studies (Banks and Blough, 2014). Second, decreasing the food reinforcer magnitude increased behavioral allocation towards methamphetamine in a concentration-dependent manner in both rat strains consistent with previous drug-vs-food choice studies in rats and monkeys (Nader and Woolverton, 1991; Negus, 2003; Thomsen et al., 2013; Townsend et al., 2021) and extending these findings to methamphetamine. Third, increasing the methamphetamine response requirement (i.e., FR) produced an FR-dependent decrease in methamphetamine choice with FR125 eliminating methamphetamine choice in both strains. Overall, these data support the hypothesis that methamphetamine and liquid food function as near-perfect substitutes in both Sprague Dawley and Long Evans rats and provide an empirical platform to determine the pharmacological mechanisms of methamphetamine reinforcement.

Two common experimental methods to assess economic substitutability between two reinforcers are to manipulate either reinforcer magnitude or reinforcer cost (i.e. response requirement). In the present study, both increasing the reinforcer magnitude for either methamphetamine or food and increasing methamphetamine cost shifted the methamphetamine-vs-food choice dose-effect function as hypothesized. The present results were consistent with the extant drug choice literature in both rats and nonhuman primates for other addictive drug classes such as cocaine (Cantin et al., 2010; Nader and Woolverton, 1991; Negus, 2003; Thomsen et al., 2013) and opioids (Doyle et al., 2023; St. Onge et al., 2022; Townsend et al., 2021). Furthermore, the preclinical results are consistent with human laboratory drug choice studies where money is the alternative nondrug reinforcer(Bennett et al., 2013; Comer et al., 1997; Heishman et al., 2000; Lile et al., 2016). The consistency of food reinforcers in preclinical studies suggest that food, regardless of food type (i.e. pellets, saccharin, sucrose, or Ensure), may function as a universal substitute for addictive drugs similar to money functioning as a universal substitute in human laboratory drug self-administration studies. Future research determining the neurobiological and behavioral mechanisms how food functions as an economic substitute for addictive drugs should improve our understanding of both general reinforcement and substance use disorders.

### 4.2 Xanomeline treatment effects on methamphetamine choice

The second aim was to determine the effectiveness of the M1/M4 agonist xanomeline to attenuate methamphetamine choice in male and female Sprague Dawley and Long Evans rats. Again, there were two main findings. First, xanomeline significantly attenuated 0.1 mg/kg/inf methamphetamine choice in Long Evans rats regardless of the xanomeline dose tested. However, xanomeline failed to attenuate methamphetamine choice in Sprague Dawley rats. Furthermore, xanomeline treatment effects were surmounted by a larger self-administered methamphetamine dose. The present results in Long Evans rats are generally consistent with published xanomeline treatment effects on cocaine choice in rats and monkeys (Marsh et al., 2025; Thomsen et al., 2014) and extend these findings to methamphetamine. Second, xanomeline treatment effects on methamphetamine choice were most robust during the first 30 min of the choice session and effectively absent during the last 30 min. The time course of xanomeline treatment effects was consistent with the short half-life of xanomeline in rats (Bymaster et al., 1997). Overall, these data suggest muscarinic receptors involvement in methamphetamine reinforcement and that xanomeline warrants further evaluation as a candidate MUD medication.

Monitoring xanomeline effects on methamphetamine choice over the 5-day treatment revealed two main findings. First, xanomeline-induced reductions in methamphetamine reinforcers were evident on Day 1 of xanomeline treatment and sustained through Day 5. These results suggest that tolerance did not develop to xanomeline treatment effects on methamphetamine self-administration supporting the conclusion that xanomeline may have clinical utility as a candidate MUD pharmacotherapy. However, clinically effective substance use disorder treatments not only decrease drug self-administration but promote behavioral reallocation towards nondrug reinforcers. Towards this second goal, xanomeline-induced behavioral reallocation away from methamphetamine and towards food was not evident on Day 1 and instead developed over the course of the 5-day treatment period. The absence of behavioral reallocation towards food at Day 1 may reflect acute muscarinic agonist undesirable effects with tolerance developing to these undesirable xanomeline effects with repeated treatment. Xanomeline is clinically available under the tradename Cobenfy ® in combination with the peripheral muscarinic antagonist trospium. Future studies could determine xanomeline treatment effects in combination with trospium to determine if the combination facilitates behavioral reallocation away from methamphetamine sooner after treatment initiation.

Future research directions involve delineating the underlying mechanisms of xanomeline treatment effectiveness on methamphetamine self-administration. In vivo microdialysis studies revealed xanomeline increased extracellular dopamine in the prefrontal cortex and the nucleus accumbens suggesting xanomeline may function as an “agonist-like” medication for methamphetamine (Perry et al., 2001) similar to treatment strategies used for opioid (methadone) and nicotine (varenicline) dependance, in addition with amphetamine maintenance for cocaine in preclinical models (Cahill et al., 2016; Mattick et al., 2009; Negus and Henningfield, 2015; Venniro et al., 2020). Lastly, the role of M1 and M4 activation for xanomeline treatment effectiveness to decrease methamphetamine self-administration remains to be fully elucidated. Knocking out the M4 receptor in mice delays acquisition of cocaine discrimination and to a lesser extent with M1 knockouts when compared training sessions to wild type(Joseph and Thomsen, 2017). A future direction of this study includes selectively targeting M1 and M4 receptors separately using the M1 bitopic agonist VU0364572 (Digby et al., 2012) and M4 positive allosteric modulator (Byun et al., 2014) to determine muscarinic receptor mechanisms of xanomeline treatment effects on methamphetamine choice.

### 4.3 Effects of rat strain on methamphetamine choice

There were no significant differences between Sprague Dawley and Long Evans rats for either reinforcer magnitude or methamphetamine cost manipulations on methamphetamine-vs-food choice. Furthermore, transforming the methamphetamine choice function to a methamphetamine demand curve also failed to reveal rat strain differences. There are a paucity of studies that systematically compared rat strain on drug self-administration endpoints. Comparing Lewis and Fischer 344 rats, food essential value was greater in Lewis compared to Fischer 344 rats, whereas cocaine essential value was the opposite with essential value greater in Fischer 344 compared to Lewis rats (Christensen et al., 2009). Future studies could use Heterogenous Stock rats to study individual and genetic factors underlying methamphetamine-vs-food choice similar to published cocaine choice studies (Sedighim et al., 2021). In contrast to the nonpharmacological manipulations, there were strain differences in xanomeline treatment effects with xanomeline only significantly attenuating methamphetamine choice in Long Evans, but not Sprague Dawley rats. Reasons for these differential xanomeline effects based on strain are unclear as previous xanomeline treatment studies (Marsh et al., 2025; Thomsen et al., 2014) on cocaine choice used Sprague Dawley rats. Furthermore, there is no published evidence on muscarinic receptor expression or distribution differences between rat strains. Moreover, these observed strain differences in xanomeline effectiveness combined with the lack of xanomeline effectiveness to attenuate 0.32 mg/kg/infusion methamphetamine choice may suggest limited clinical utility of xanomeline as a candidate MUD pharmacotherapy compared to cocaine use disorder.

## Acknowledgements and Funding

Research was supported by the National Institute of Health grants R01DA055825 and T32GM148403. The National Institute on Drug Abuse had no role in study design, collection, analysis or interpretation of the data, in the writing or decision to submit the manuscript for publication. The manuscript content is solely the responsibility of the author and does not necessarily reflect the official views of the National Institutes of Health.

## Author Contributions

Amber Baldwin was responsible for original conceptualization, methodology, investigation, writing the original draft, reviewing, and editing. Matthew Banks was responsible for original conceptualization, refining methodology, funding acquisition, project administration, supervision, reviewing, and editing.

## References

1. Acuff, S.F., MacKillop, J., Murphy, J.G., 2023. A contextualized reinforcer pathology approach to addiction. Nat Rev Psychol 2(5), 309–323.

2. Banks, M., Blough, B., 2014. Effects of Pharmacological and Environmental Manipulations on Methamphetamine vs. Food Choice in Rhesus Monkeys. Neuropsychopharmacology 39, S591.

3. Banks, M.L., Blough, B.E., 2015. Effects of environmental maniuplations and bupropion and risperidone treatments on choice between methamphetamine and food in rhesus monkeys. Neuropsychoharmacology 40(9), 2198–2206.

4. Banks, M.L., Negus, S.S., 2017. Insights from Preclinical Choice Models on Treating Drug Addiction. Trends Pharmacol Sci 38, 181–194.

5. Bennett, J.A., Stoops, W.W., Rush, C.R., 2013. Alternative reinforcer response cost impacts methamphetamine choice in humans. Pharmacol Biochem Behav 103(3), 481–486.

6. Bickel, W.K., DeGrandpre, R.J., Higgins, S.T., 1995. The behavioral economics of concurrent drug reinforcers: a review and reanalysis of drug self-administration research. Psychopharmacology 118(3), 250–259.

7. Bymaster, F.P., Whitesitt, C.A., Shannon, H.E., DeLapp, N., Ward, J.S., Calligaro, D.O., Shipley, L.A., Buelke-Sam, J.L., Bodick, N.C., Farde, L., Sheardown, M.J., Olesen, P.H., Hansen, K.T., Suzdak, P.D., Swedberg, M.D.B., Sauerberg, P., Mitch, C.H., 1997. Xanomeline: A selective muscarinic agonist for the treatment of Alzheimer’s disease. Drug Development Research 40(2), 158–170.

8. Byun, N.E., Grannan, M., Bubser, M., Barry, R.L., Thompson, A., Rosanelli, J., Gowrishankar, R., Kelm, N.D., Damon, S., Bridges, T.M., Melancon, B.J., Tarr, J.C., Brogan, J.T., Avison, M.J., Deutch, A.Y., Wess, J., Wood, M.R., Lindsley, C.W., Gore, J.C., Conn, P.J., Jones, C.K., 2014. Antipsychotic Drug-Like Effects of the Selective M4 Muscarinic Acetylcholine Receptor Positive Allosteric Modulator VU0152100. Neuropsychopharmacology 39(7), 1578–1593.

9. Cahill, K., Lindson-Hawley, N., Thomas, K.H., Fanshawe, T.R., Lancaster, T., 2016. Nicotine receptor partial agonists for smoking cessation. Cochrane Database of Syst Rev (5).

10. Cantin, L., Lenoir, M., Augier, E., Vanhille, N., Dubreucq, S., Serre, F., Vouillac, C., Ahmed, S.H., 2010. Cocaine Is Low on the Value Ladder of Rats: Possible Evidence for Resilience to Addiction. PLoS ONE 5(7), e11592.

11. Christensen, C.J., Kohut, S.J., Handler, S., Silberberg, A., Riley, A.L., 2009. Demand for food and cocaine in Fischer and Lewis rats. Behav Neurosci 123(1), 165–171.

12. Comer, S.D., Collins, E.D., Fischman, M.W., 1997. Choice between money and intranasal heroin in morphine-maintained humans. Behav Pharmacol 8(8), 677–690.

13. Digby, G.J., Noetzel, M.J., Bubser, M., Utley, T.J., Walker, A.G., Byun, N.E., Lebois, E.P., Xiang, Z., Sheffler, D.J., Cho, H.P., Davis, A.A., Nemirovsky, N.E., Mennenga, S.E., Camp, B.W., Bimonte-Nelson, H.A., Bode, J., Italiano, K., Morrison, R., Daniels, J.S., Niswender, C.M., Olive, M.F., Lindsley, C.W., Jones, C.K., Conn, P.J., 2012. Novel Allosteric Agonists of M_1_ Muscarinic Acetylcholine Receptors Induce Brain Region-Specific Responses That Correspond with Behavioral Effects in Animal Models. The Journal of Neuroscience 32(25), 8532–8544.

14. Doyle, W.S., Freeman, K.B., Woods, J., Huskinson, S.L., 2023. Choice between food and cocaine or fentanyl reinforcers under fixed and variable schedules in female and male rhesus monkeys. Psychopharmacology 240(7), 1573–1585.

15. Fischer, B., O’Keefe-Markman, C., Lee, A., Daldegan-Bueno, D., 2021. ‘Resurgent’, ‘twin’ or ‘silent’ epidemic? A select data overview and observations on increasing psycho-stimulant use and harms in North America. Substance Abuse Treatment, Prevention, and Policy 16(1), 17.

16. Heishman, S.J., Schuh, K.J., Schuster, C.R., Henningfield, J.E., Goldberg, S.R., 2000. Reinforcing and subjective effects of morphine in human opioid abusers: effect of dose and alternative reinforcer. Psychopharmacology 148(3), 272–280.

17. Hersch, S., Gutekunst, C., Rees, H., Heilman, C., Levey, A., 1994. Distribution of m1-m4 muscarinic receptor proteins in the rat striatum: light and electron microscopic immunocytochemistry using subtype-specific antibodies. The Journal of Neuroscience 14(5), 3351–3363.

18. Holly, E.N., Galanaugh, J., Fuccillo, M.V., 2024. Local regulation of striatal dopamine: A diversity of circuit mechanisms for a diversity of behavioral functions? Current Opinion in Neurobiology 85, 102839.

19. Hursh, S.R., Silberberg, A., 2008. Economic demand and essential value. Psychological Review 115(1), 186–198.

20. Joseph, L., Thomsen, M., 2017. Effects of muscarinic receptor antagonists on cocaine discrimination in wild-type mice and in muscarinic receptor M1, M2, and M4 receptor knockout mice. Behavioural Brain Research 329, 75–83.

21. Kearns, D.N., 2025. A Review of Behavioral Economic Manipulations Affecting Drug versus Nondrug Choice in Rats. Perspect Behav Sci 48(2), 341–366.

22. Kearns, D.N., Kim, J.S., Tunstall, B.J., Silberberg, A., 2016. Essential values of cocaine and non-drug alternatives predict the choice between them. Addict Biol, n/a–n/a.

23. Lamb, R.J., Ginsburg, B.C., 2017. Addiction as a BAD, a Behavioral Allocation Disorder. Pharmacology Biochemistry and Behavior.

24. Lile, J.A., Stoops, W.W., Rush, C.R., Negus, S.S., Glaser, P.E.A., Hatton, K.W., Hays, L.R., 2016. Development of a translational model to screen medications for cocaine use disorder II: Choice between intravenous cocaine and money in humans. Drug Alcohol Depend.

25. Marcus, M.M., Banks, M.L., 2025. Mechanistic and translational insights from preclinical cocaine choice procedures on the economic substitutability of cocaine and nondrug reinforcers. Neuroscience & Biobehavioral Reviews 175, 106217.

26. Marsh, S.A., Heslep, N., Paronis, C.A., Bergman, J., Negus, S.S., Banks, M.L., 2025. Xanomeline treatment attenuates cocaine self-administration in rats and nonhuman primates. Neuropharmacology 281, 110686.

27. Mattick, R.P., Breen, C., Kimber, J., Davoli, M., 2009. Methadone maintenance therapy versus no opioid replacement therapy for opioid dependence. Cochrane Database of Systematic Reviews (3).

28. Mirza, N.R., Peters, D., Sparks, R.G., 2003. Xanomeline and the Antipsychotic Potential of Muscarinic Receptor Subtype Selective Agonists. CNS Drug Reviews 9(2), 159–186.

29. Nader, M.A., Woolverton, W.L., 1991. Effects of increasing the magnitude of an alternative reinforcer on drug choice in a discrete-trials choice procedure. Psychopharmacology 105(2), 169–174.

30. Negus, S.S., 2003. Rapid Assessment of Choice between Cocaine and Food in Rhesus Monkeys: Effects of Environmental Manipulations and Treatment with d-Amphetamine and Flupenthixol. Neuropsychopharmacology 28(5), 919–931.

31. Negus, S.S., Henningfield, J., 2015. Agonist Medications for the Treatment of Cocaine Use Disorder. Neuropsychopharmacology 40(8), 1815–1825.

32. Perry, K.W., Nisenbaum, L.K., George, C.A., Shannon, H.E., Felder, C.C., Bymaster, F.P., 2001. The muscarinic agonist xanomeline increases monoamine release and immediate early gene expression in the rat prefrontal cortex. Biological Psychiatry 49(8), 716–725.

33. Russo, S.J., Nestler, E.J., 2013. The brain reward circuitry in mood disorders. Nature Reviews Neuroscience 14(9), 609–625.

34. Sedighim, S., Carrette, L.L.G., Venniro, M., Shaham, Y., de Guglielmo, G., George, O., 2021. Individual differences in addiction-like behaviors and choice between cocaine versus food in Heterogeneous Stock rats. Psychopharmacology 238(12), 3423–3433.

35. St. Onge, C.M., Canfield, J.R., Ortiz, A., Sprague, J.E., Banks, M.L., 2024. Xylazine does not enhance fentanyl reinforcement in rats: A behavioral economic analysis. Drug and Alcohol Dependence 258, 111282.

36. St. Onge, C.M., Taylor, K.M., Marcus, M.M., Townsend, E.A., 2022. Sensitivity of a fentanyl-vs.-social interaction choice procedure to environmental and pharmacological manipulations. Pharmacology Biochemistry and Behavior 221, 173473.

37. Stocco, M.R., Purpura, M., Vieira, P.A., Wallquist, K., Wang, S., Adams, J., Szumlinski, K.K., Kippin, T.E., 2025. Time to choose: impact of intertrial interval on selecting between methamphetamine and food reinforcement in male and female rats. Psychopharmacology 242(4), 693–702.

38. Swerdlow, N.R., Krupin, A.S., Bongiovanni, M.J., Shoemaker, J.M., Goins, J.C., Hammer, R.P., 2006. Heritable Differences in the Dopaminergic Regulation of Behavior in Rats: Relationship to D2-Like Receptor G-Protein Function. Neuropsychopharmacology 31(4), 721–729.

39. System, N.F.L.I., 2023. NFLIS Drug Snapshot, in: Justice, D.o. (Ed.).

40. Thomsen, M., Barrett, A.C., Negus, S.S., Caine, S.B., 2013. Cocaine versus food choice procedure in rats: environmental manipulations and effects of amphetamine. Journal of the experimental analysis of behavior 99(2), 211–233.

41. Thomsen, M., Fulton, B., Caine, S.B., 2014. Acute and chronic effects of the M1/M4-preferring muscarinic agonist xanomeline on cocaine vs. food choice in rats. Psychopharmacology 231(3), 469–479.

42. Townsend, E.A., Negus, S.S., Caine, S.B., Thomsen, M., Banks, M.L., 2019. Sex differences in opioid reinforcement under a fentanyl vs. food choice procedure in rats. Neuropsychopharmacology.

43. Townsend, E.A., Schwienteck, K.L., Robinson, H.L., Lawson, S.T., Banks, M.L., 2021. A drug-vs-food “choice” self-administration procedure in rats to investigate pharmacological and environmental mechanisms of substance use disorders. Journal of Neuroscience Methods, 109110.

44. Venniro, M., Banks, M.L., Heilig, M., Epstein, D.H., Shaham, Y., 2020. Improving translation of animal models of addiction and relapse by reverse translation. Nature Reviews Neuroscience.

45. Venniro, M., Zhang, M., Caprioli, D., Hoots, J.K., Golden, S.A., Heins, C., Morales, M., Epstein, D.H., Shaham, Y., 2018. Volitional social interaction prevents drug addiction in rat models. Nature Neuroscience 21(11), 1520–1529.

46. Weikop, P., Jensen, K.L., Thomsen, M., 2020. Effects of muscarinic M1 receptor stimulation on reinforcing and neurochemical effects of cocaine in rats. Neuropsychopharmacology 45(12), 1994–2002.

47. Weiner, D.M., Levey, A.I., Brann, M.R., 1990. Expression of muscarinic acetylcholine and dopamine receptor mRNAs in rat basal ganglia. Proceedings of the National Academy of Sciences 87(18), 7050–7054.

